# Uncovering the principles coordinating systems-level organelle biogenesis with cellular growth

**DOI:** 10.1101/2022.11.01.514705

**Authors:** Shixing Wang, Shankar Mukherji

## Abstract

Among the hallmark properties of the eukaryotic cell is its organization into specialized biochemical compartments known as organelles. Understanding how organelle biogenesis at systems-scale is coordinated with cellular growth rate and size is a major goal of quantitative cell biology. Here we map out the correlation structure of systems-level organelle biogenesis with cellular growth using “rainbow yeast”, a strain of *Saccharomyces cerevisiae* that expresses fluorescent labels for 6 major organelles. By carrying out hyperspectral imaging of thousands of single rainbow yeast cells, we decomposed the systems-level organelle biogenesis program into specific modes that characterize the response to changes in nutrient availability. Upon chemical biological dissection of this response, our results suggest that systems-level organelle biogenesis represents the sum of distinct organelle modes excited by growth rate and cell size separately. The flexibility afforded by this regulatory architecture may underlie how eukaryotic cells leverage compartmentalization to independently tune cell sizes and growth rates and satisfy potentially incompatible environmental and developmental constraints.

## Main Text

The principles by which cells coordinate organelle growth with overall cellular growth have been examined primarily through scaling relationships of organelle sizes with the sizes of their host cells (*1–10*). It is widely appreciated, for example, that the volume of the nucleus scales linearly with the size of its host cell (*1–4*). Similar scaling relationships have been documented for the endoplasmic reticulum (*5*), vacuoles/lysosomes (*6,7*), and mitochondria (*8,9*). However our knowledge of pairwise scaling relationships among organelles is sparse. Even less characterized are multi-organelle interactions known to be important for regulating cellular scale physiology (*10*). We reasoned that a combination of systems-scale organelle imaging inspired by previous efforts (*11*,*12*), chemical biology tools (*13–15*), and simple data reduction methods would yield deep insights into how organelle biogenesis couples to the interrelated variables of cell growth rate and cell size.

In order to visualize the 6 major organelles in budding yeast, we transcriptionally fused genes encoding fluorescent proteins to genes encoding organelle-resident proteins known with high confidence to localize to a specific organelle (*16*, Fig. 1A,B, Table S1). Rainbow yeast cells were then cultured in defined media containing varying amounts of glucose, logarithmically spaced from 0% to 2%, and subsequently imaged using a confocal laser-scanning microscope equipped with a diffraction grating that resolved photons emitted from our sample onto a multianode photomultiplier tube. The subsequent pixel-by-pixel spectra were then unmixed using experimentally derived spectra of the fluorescent proteins resident to each organelle, resulting in images in which each cell contained 6 image channels, one corresponding to each organelle under study (Fig. S1). Machine learning-assisted segmentation of yeast cell boundaries and individual organelle images (*16–18,* Fig. S2) resulted in a dataset in which each cell was associated with the organelle properties from that cell, such as organelle number, the total organelle volume, and the fraction of the cell volume taken up by the summed volumes of a given organelle type. While the measured organelle sizes and volume fractions are potentially overestimates due to limitations from diffraction, our measurements have sufficient resolving power to detect differences in these properties from one cell to another despite the systematic error.

**Fig. 1:**
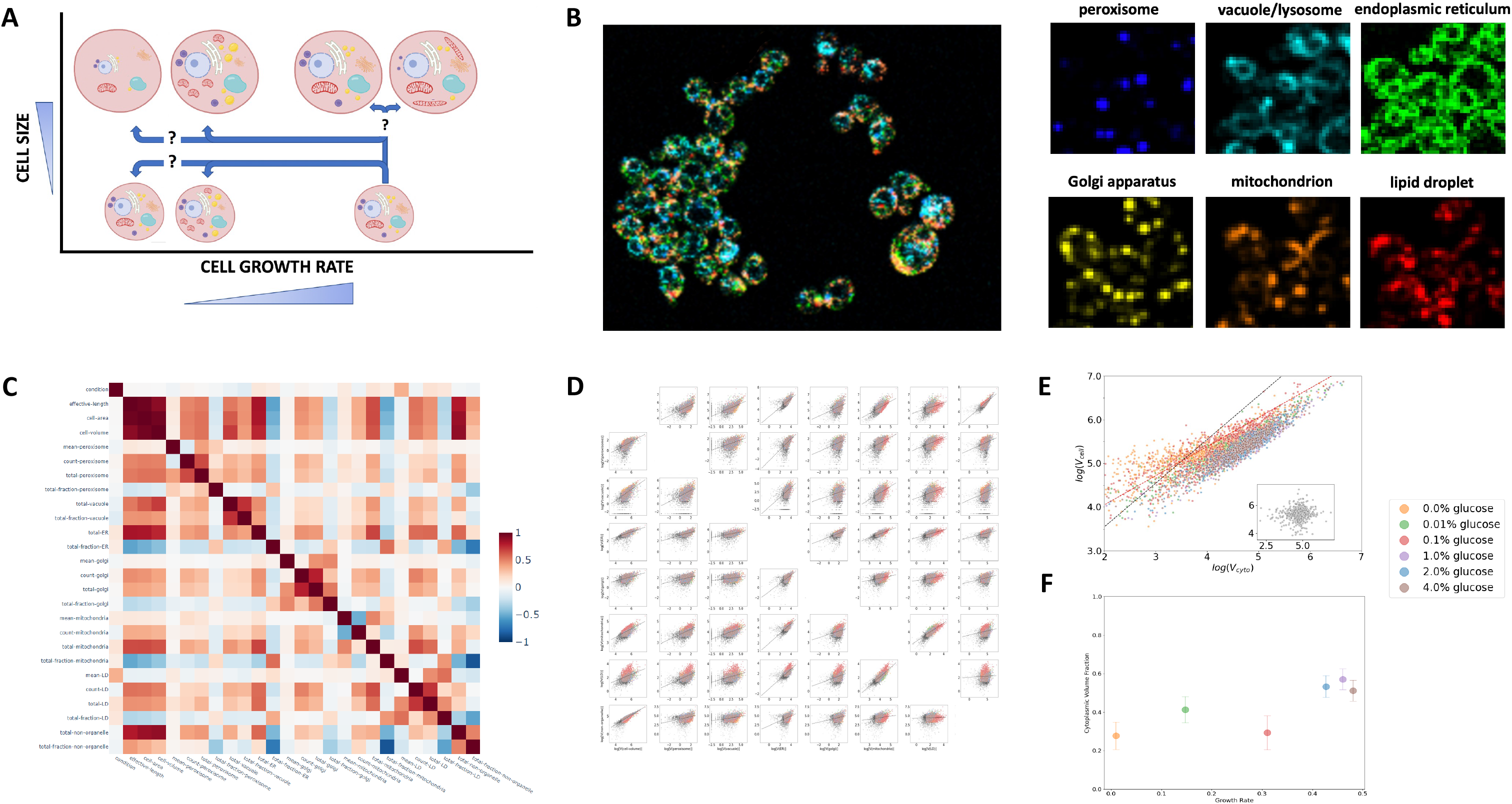
**(A)**Schematic depicting possible responses of systems-level organelle biogenesis upon variations in cellular size and growth rate. **(B)**False color images of individual rainbow yeast cells following linear unmixing of image data obtained from hyperspectral laser-scanning confocal microscopy. Left panel: overlay of all 6 organelles. Right panels: individual channels of organelles (blue: peroxisome, cyan: vacuole, green: endoplasmic reticulum, yellow: Golgi apparatus, orange: mitochondrion, red: lipid droplet). **(C)**Heatmap of correlation coefficients between organelle and cellular properties of each cell upon glucose perturbation. **(D)**Log-log plots of total volumes of 6 organelles, total volume of the cell, and estimate of nucleocytoplasmic volume. The diagonal plots are the histograms of logarithm of organelles’ total volumes. Lines are the linear regression within each condition. **(E)**Log-log scatter plot of the cellular volume versus the cytoplasmic volume. **(F)**Scatter plot of the nucleocytoplasmic volume fraction versus cellular growth rate under different glucose concentrations. The error bars show the standard error around the mean.

In order to validate our data, we generated a pairwise correlation coefficient matrix between the different organelle properties we examined and compared the entries in this matrix to previous studies (Fig. 1C). In each case we examined we found good agreement between the results of our systems-scale organelle imaging and previous results. For example, examining the relationship between total organelle size and the size of its host cells, we recover the positive correlation observed previously for the ER, vacuole/lysosome, and mitochondria (Fig. 1C).

With a validated dataset in hand, we sought to discriminate between various models that aim to capture the dominant constraints from cellular compartments on cell size and growth. To this end, we extracted volume scaling exponents between every pair of structures we imaged as well as the volume of the cell and an estimate of the volume of the nucleocytoplasm (Fig. 1D, Table S2). The exponents revealed a variety of relationships, ranging from relatively insensitive scaling of the vacuole to the other organelles in the cell, to isometric scaling of endomembrane organelles such as the Golgi, peroxisome, and lipid droplet with the ER. In particular, we sought to estimate how the volume of the cell scales with the volume of the nucleocytoplasm. Theoretical predictions of the scaling exponent are marked by disagreements depending on the process proposed to limit cell size: limitations on cell size from bulk protein synthesis tend to favor exponents of 1, while transport limited cell size regulation tend to favor exponents of 2/3. We observed that for each glucose concentration, the volume of the cell obeyed a scaling exponent of 0.56 +/− 0.01 (Fig. 1E). Given the strong correlation between cell size and growth rate, the sublinear scaling exponent predicts that the volume fraction of the cell devoted to nucleocytoplasm should increase with increasing growth rate, which we observe (Fig. 1F).

The decreasing availability of cellular volume for organelles due to demand from increasing nucleocytoplasmic volume immediately raised the question of whether all organelles are equally scaled down to allow nucleocytoplasmic growth or whether certain organelles are prioritized over others upon changes in cell size and growth rate. To address these questions we carried out two analyses.

First, in order to quantify how organized changes in systems-level organelle biogenesis are, we analyzed the coherence with which the organelle profiles of the cells change upon changes in glucose. In order to capture how the cell allocates its limited material and energy budget, we focused our analysis on organelle volume fractions. Using the pixel-level posterior probabilities of assigning a given pixel to a given type of organelle, we estimate that the errors in our volume fraction measurements range from 5-10% (*16*, Fig. S3). To measure the change, we computed the Kullback-Leibler divergence between organelle volume fractions distributions from populations of cells grown at different glucose concentrations, using 2% glucose as the standard reference concentration to measure with respect to. To measure the coherence of any change, we computed the Shannon entropy of the organelle volume fraction distributions of each population. By comparing the relative magnitude of changes between the Shannon entropy and KL divergence as cells were grown in different glucose concentrations, we observed that the KL divergence increased by 200-400% compared to the reference population, indicating that cells do alter their organelle allocations in response to changes in glucose, but the Shannon entropy changed by 10% (Fig. S4). This suggests that the organelle profile of cells are equally constrained at low and high glucose levels.

As our results suggested that cells follow a relatively stereotyped set of changes in organelle volume fractions, we sought to identify these changes. To this end, we subjected the results of our hyperspectral imaging of rainbow yeast to principal components analysis and compute a vector, which we term the condition vector, that connects the centroids of the data from each experimental condition (Fig. 2A, *19*). Upon examination of the first 3 principal components, we see a strong relationship between a single cell’s position in this space and the glucose environment it was cultured in (Fig. 2B, S5). In order to gain cell biological insight from the structure of the data along its principal components, we plot the principal components in a basis space spanned by unit vectors in the directions of most significant organelle volume fractions (Fig. 2C) and compute how co-linear these principal components are with the condition vector averaged over multiple glucose conditions. We see that in this basis, the first and second principal components consist of decreasing organelle volume fractions upon increase in environmental glucose concentration (Fig. 2C,D). The third principal component, however, is dominated by a combination of decreasing mitochondrial and lipid droplet volume fractions concomitant with increasing peroxisomal and Golgi volume fractions (Fig. 2C,D). Rather than acting independently, our systems-level view uncovers patterns of correlated organelle biogenesis in response to glucose availability. We interpret this data to mean that, in agreement with what would be expected from previous knowledge about yeast physiology in which decreasing glucose concentrations lead to increasing contributions from respiration over fermentation, individual cells preferentially downregulate their mitochondrial and lipid droplet volume fractions upon increasing levels of glucose while elevating resources devoted to their peroxisomes and Golgi.

**Fig. 2:**
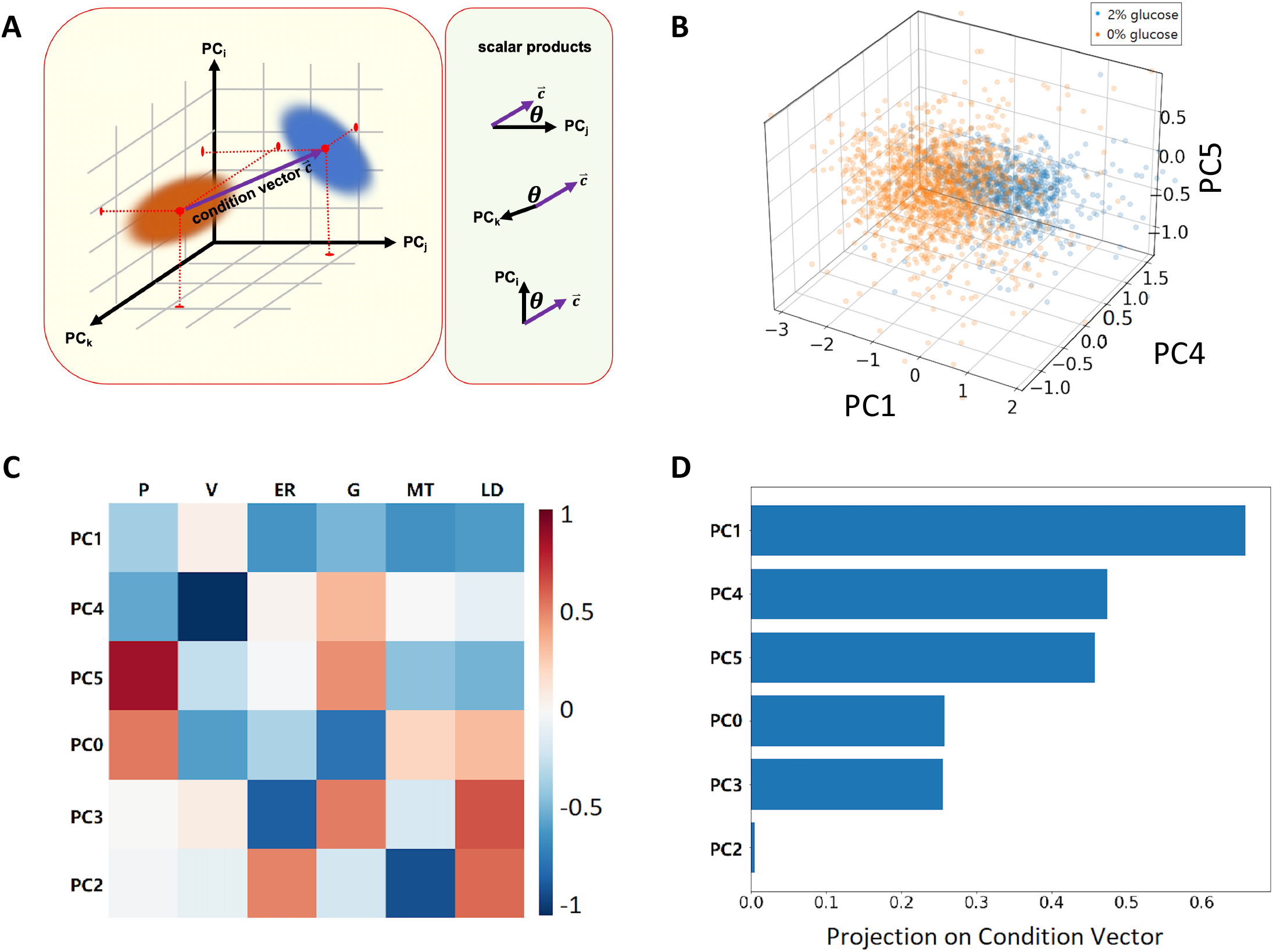
**(A)**Schematic depiction of the condition vector in the PCA space. **(B)**Projection of normalized organelle volume fraction onto the subspace extended by 3 principal components whose direction are closest to the condition vector. **(C)**Heatmap of principal components of normalized organelle volume fractions. Rows are sorted in descending order of the inner products of the principal components and the condition vector, and columns are the organelles (P: peroxisome, V: vacuole, ER: endoplasmic reticulum, G: Golgi apparatus, MT: mitochondrion, LD: lipid droplet). **(D)**Cosine of the angle between the condition vector and different principal components, sorted in descending order.

Our results, however, raise a fundamental question regarding the rules by which the cell organizes systems-level organelle biogenesis: does the organelle biogenesis program track the size of the cell alone or growth rate of the cell (Fig. 1A)? Limitation of nutrients as experienced in natural settings, such as in the case of glucose, typically results in both cell size and growth rate reductions (*20–26*). In order to distinguish these possibilities we compared the results of two further analyses of our rainbow yeast cells. To examine the effects of nutrient-indepdendent cell size control, we transformed a plasmid containing a copy of the gene Whi5 under control of a *β*-estradiol inducible promoter (*27,28*). While the mechanism by which Whi5 expression levels alter cell size are under investigation, in our experiments we observe a roughly 2.4-fold increase in cell size upon exposure of cells to saturating levels of *β*-estradiol. To examine the effects of growth rate dependence, we leveraged the fact that rainbow yeast was built on a genetic background that renders the cells auxotrophic for the amino acid leucine. We could then titrate leucine levels in the growth medium and thereby affect growth rate (*29*). We then performed hyperspectral imaging on rainbow yeast cells at various leucine and *β*-estradiol concentrations present in the culture medium.

By examining the scalar products between the principal components obtained from cells grown in varying glucose concentrations and the principal components obtained from cells grown in either varying leucine (Fig. 3A,B, S6) or varying *β*-estradiol concentrations (Fig. 3D,E), we can dissect the systems-level organelle biogenesis response to glucose into growth rate-responsive and cell size responsive organelle modes. In the case of cell size, systems-level organelle volume fraction modes are dominated by variation along the first principal component, which consists of decreasing organelle volume fractions (Fig. 3B,C). Leucine limitation, on the other hand, results in organelle modes dominated by a principal component in which mitochondrial and lipid droplet volume fractions decrease while Golgi and peroxisome volume fractions increase upon increasing leucine concentration (Fig. 3E,F). The data thus suggests that the signal dominating systems-level organelle prioritization comes from the altered growth rate rather than cell size alone.

**Fig. 3:**
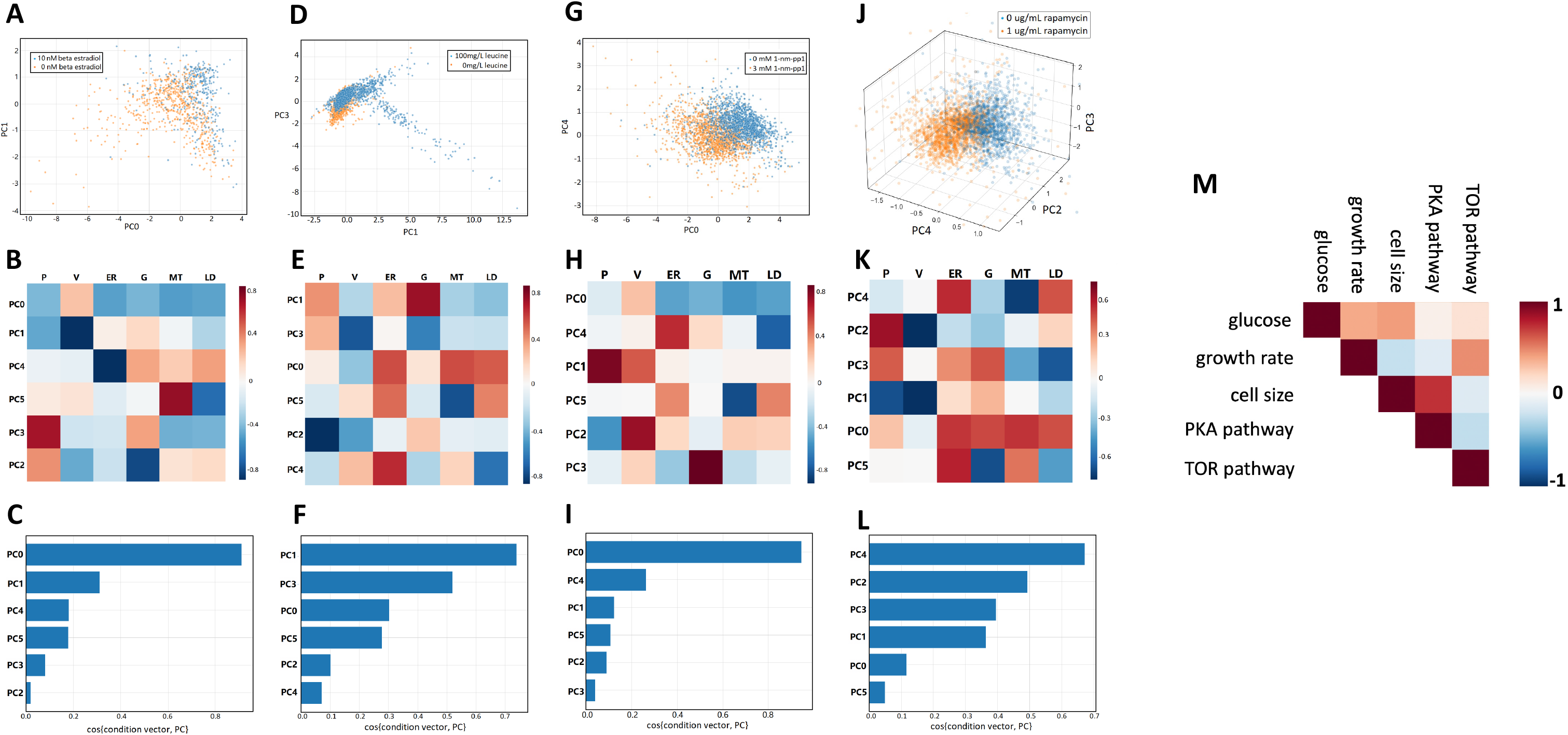
**(A,D,G,J)**Projection of normalized organelle volume fraction onto the subspace extended by principal components whose direction are closest to the condition vector in different experiments (**A**: Whi5 overexpression strain treated by *β*-estradiol, **D**: leucine, **G**: 1nm-pp1, **J**: rapamycin). **(B,E,H,K)**Heatmap of principal components of normalized organelle volume fractions, sorted by the inner product of the principal components and the condition vector. Plots are arranged by experiments in the same order as PCA projections. **(C,F,I,L)**Cosine of the angle between the condition vector and different principal components across different experiments. Plots are arranged in the same order as PCA projections. **(M)**Pairwise similarities of principal components across different experiments. The similarity is calculated by averaging the inner products of each pair of principal components in two experiments, weighted by the cosine of the angles between the principal components and the experiment’s condition vector.

Finally, as organelles represent a significant cellular expense in terms of material and energy, we aimed to use our measurement and analysis strategy to gain mechanistic insight into whether systems-scale organelle biogenesis are driven by physical constraints from cell size or nutrient availability or whether information from growth regulating signaling pathways alone could provoke the changes observed in otherwise unperturbed cells. The two principal growth signaling pathways in budding yeast are the target-of-rapamycin (TOR) and protein kinase A (PKA) pathways (*30,31*). We sought to characterize the systems-level organelle biogenesis profile in the basis of organelle fractions as a function of TOR, focusing on TORC1 activity, and PKA signal strengths. While in the case of the TOR pathway this is possible to accomplish using the small molecule inhibitor rapamycin, in the case of the PKA pathway we created rainbow yeast in a genetic background that carries mutations in the PKA pathway genes TPK1, TPK2, and TPK3 that render these enzymes inhibitable by the small molecule 1-nm-pp1 (*13*). We then subjected rainbow yeast to varying concentrations of rapamycin and 1-nm-pp1, performed hyperspectral imaging, and plotted the principal components in the organelle volume fraction basis space (Fig. 3G, S7). We observe that the first principal component in the case of TOR signaling inhibition tracks closely to the organelle modes responsive to cell growth rate (Fig. 3G–I, M). On the other hand, the organelle mode principally excited by PKA pathway (Fig. 3J–L, S8) inhibition is largely co-linear with the mode responsive to cell size (Fig. 3M). This data suggests that TOR and PKA signaling link cell growth rate and cell size respectively to systems-level organelle biogenesis.

The highly interactive yet modular nature of organelles and the complex, multiscale processes that govern their function make elucidating the principles by which their biogenesis is regulated difficult (*32*). By employing a top-down, data-driven approach we suggest that organelle biogenesis in response to physiological cues couples organelles to each other in stereotyped ways. Our map of pairwise correlations revealed that organelle volume fractions generally anti-correlate with host cell size. This observation suggests that, by growing organelles with size scaling exponents less than 1, a greater fraction of the cell may be devoted to ribosomes even while increasing total organelle volumes needed to service faster growth (*33,34*). Our data also suggests that the coordination of systems-level organelle biogenesis with the host cell is achieved by coupling their growth rates rather than solely allocating fixed fractions of their cell size. Our data suggests that PKA and TOR signaling activity mechanistically links growth rate and cell size separately to organelle biogenesis, a conclusion that would be difficult to draw without our systems-level view. Our measurement strategy, however, is highly generalizable, for example to capture how systems-level organelle biogenesis responds to diverse physiological states, such as those seen in aging cells, or in complex multicellular tissues, where organelle biogenesis must simultaneously satisfy both universal cell biological and cell-type specific metabolic demands.

## Acknowledgements

We thank M. Tikhonov for discussions related to this work. We thank K. Amiri, A. Arra, L. Perez, R. Dale, A. Singh, D. Kailash, and R. Hanley for critical feedback on the manuscript. Plasmids encoding mTagBFP2-PTS1, superfolderGFP, yemCitrine, tdTomato, and Erg6-mCherry were obtained from Addgene.

