## Supplementary Information for "Uncovering the principles coordinating systems-level organelle biogenesis with cellular growth"

**1. Construction of multi-organelle fluorescent protein labeled yeast strain.** To construct the diploid strain of *Saccharomyces cerevisiae* that tagged 6 organelles, the 2 haploid strains (EY2795 and EY2796) were mated together on a selection plate, after each consecutively tagged 3 organelles by inserting each gene of fluorescent protein before the stop codon of the gene localized to the given organelle. Table S1 shows for each organelle, the corresponding template plasmid containing the sequence of fluorescent protein and selection marker, the gene expressed locally in the organelle, and the haploid strain that the fluorescent protein gene got transformed into. mCherry-TRP and mtagBFP2-TRP were integrated into the yeast genome by linearizing and transforming the respective plasmids into the yeast. The other fluorescent protein genes were obtained with PCR and the products were transformed into the yeasts. For each PCR, a pair of forward and reverse sequence-specific primers were synthesized. The forward primers contains 40 bases upstream to the stop codon of organelle-specific gene and the beginning 20 bases of the fluorescent protein gene, while the reverse primer contains the reversed complementary sequence of 20 bases of the stop codon of the marker on the template plasmid and 40 base pairs downstream the stop codon of the organelle-specific gene. The transformed cells were selected on dropout or antibiotic plates according to the selection markers. mTFP1 sequence were custom synthesized and inserted into a PKT backbone with a KAN marker.

| Haploid | Organelle | Localized Gene | Fluorescent Protein | Selection Marker | Template Plasmid | Addgene # |
|---|---|---|---|---|---|---|
| EY2795 | vacuole | VPH1 | mTFP1 | KAN | customized PKT | n/a |
| EY2795 | Golgi apparatus | SEC7 | mCitrine | HIS | pKT0211(Sheff & Thorn 2004) | #8734 |
| EY2795 | lipid droplet | ERG6 | mCherry | TRP | ZJOM70(Zhu et al. 2019) | #133658 |
| EY2796 | peroxisome | C-SKL | mtagBFP2 | TRP | ZJOM160(Zhu et al. 2019) | #133670 |
| EY2796 | endoplasmic reticulum | SEC61 | superfolderGFP | URA | pFA6a-link-yoSuperfolderGFP-CaURA3(Lee et al. 2013) | #44873 |
| EY2796 | mitochondrion | TOM70 | tdTomato | HIS | pFA6a-link-tdTomato-SpHis5(Lee et al. 2013) | #44640 |

Table S1. Fluorescent proteins used to visualize organelles in rainbow yeast.

**2. Culturing Conditions.** Table S2 shows the different media used in the experiments. On the evening before every experiment, the yeast cells were transferred from the petri dish to 10 mL SD complete media, and grown overnight in a thermostatic shaker. For the Whi5 overexpression experiment, 10 $\mu$M of beta estradiol was added to the experimental group, and the cells were grown in the shaker overnight before microscopy imaging. For experiments other than the Whi5 overexpression, on the next morning the overnight media was diluted to OD(600nm)=0.1 in SD complete media, and grown in the shaker for 2 hours to resume exponential growth. Then the cells were washed twice and transferred into the experimental media. After 3 hours of growth in the experimental media, the cells were harvested and observed under the microscope.

* Department of Physics, Washington University in St. Louis.

| Experiment | Base Medium | Change/Addition | Condition | Alias Name | N (cells) |
|---|---|---|---|---|---|
| glucose | YNB+CSM | glucose | 0 | glu-0 | 1520 |
| | | | 0.01% m/v | glu-0-5 | 868 |
| | | | 0.1% m/v | glu-5 | 1191 |
| | | | 1% m/v | glu-50 | 555 |
| | | | 2% m/v | glu-100 | 861 |
| | | | 4% m/v | glu-200 | 790 |
| leucine | SD-leucine | leucine | 0 | leu-0 | 1012 |
| | | | 25 mg/L | leu-25 | 736 |
| | | | 50 mg/L | leu-50 | 1666 |
| | | | 75 mg/L | leu-75 | 1080 |
| | | | 100 mg/L | leu-100 | 1505 |
| cell size | SD complete | Whi5 overexpression strain | 0 | Whi5Up-betaEstradiol-0 | 651 |
| | | beta-estradiol | 10 $\mu$M | Whi5Up-betaEstradiol-10 | 353 |
| PKA pathway | SD complete | 1-nm-pp1 | 0 | 1nmpp1-0 | 1640 |
| | | | 500 nM | 1nmpp1-500 | 1554 |
| | | | 1.5 $\mu$M | 1nmpp1-1500 | 1173 |
| | | | 3 $\mu$M | 1nmpp1-3000 | 1682 |
| TOR pathway | SD complete | rapamycin | 0 | rpmc-0 | 1278 |
| | | | 100 ng/mL | rpmc-100 | 1297 |
| | | | 200 ng/mL | rpmc-200 | 772 |
| | | | 400 ng/mL | rpmc-400 | 1760 |
| | | | 1000 ng/mL | rpmc-1000 | 1963 |

Table S2. Culturing media in experiments and the corresponding alias names in dataset.

| Organelles | Excitation wavelength | Detecting wavelength | Spectral resolution |
|---|---|---|---|
| peroxisome & vacuole | 445 nm | 460 nm - 510 nm | 10 nm |
| endoplasmic reticulum | 488 nm | 500 nm - 550 nm | 50 nm |
| Golgi apparatus | 514 nm | 520 nm - 558 nm | 38 nm |
| mitochondrion & lipid droplet | 561 nm | 567 nm - 647 nm | 10 nm |

Table S3. Optical configuration of the image acquisition

**3. Image Acquisition.** The microscope slides were made from harvested cells in the experimental media and observed under a Nikon Ti2 microscope. Image acquisition started after the cells stopped moving in the field of view. A bright field microscopy image of the focal plane was taken first. Then z-stack images of hyper-spectral confocal microscopy at 4 different optical configurations were taken. (See Table 3) After that, another single-z bright field microscopy image was taken slightly off focus in order to segment the cells (see image analysis section).

**4. Image Processing.** Other than the third-party tools specified below, we used scikit-image (van der Walt et al. 2014), a python image processing package.

To segment the cells, YeaZ (Dietler et al. 2020), a convolutional neural network for yeast cellular segmentation was used. The bright field microscopy images before and after the hyper-spectral imaging were registered from the bright field camera to the hyper-spectral detector, and applied a histogram equalization filter, and then were fed into YeaZ. The resulting segmentations from the before- and after- images were compared manually, and the cells that moved during the image acquisition were excluded. In the experiment of leucine, bright field images were not taken, and the segmentation was performed on the average z-projection of the green channel hyperspectral confocal microscopy images, by manual thresholding and watershed with ImageJ (Schindelin et al. 2012).

To get single-channel z-stack images of individual organelles, the blue and red channels of hyper-spectral confocal microscopy images were demixed with the Nikon Ti2 Analysis software, by manually choosing the regions of interest for the software to extract the spectral characteristics. To segment the organelles, ilastik, an interactive machine learning software (Berg et al. 2019) was used on the demixed single-organelle images. Before feeding into ilastik, the yellow and red channel images were cleaned by subtracting the average pixel intensity of the neighborhood around each cell. Then a 3D Gaussian filter with sigma=0.75 was applied to all images. For each organelle, an ilastik project was trained with hand-drawn foreground-background labels on a sample image.

The binary images that ilastik produces were post-processed according to the type of organelles they contained. For small and globular organelles such as the peroxisome, the Golgi apparatus, and the lipid droplet, the binary images were watershed using the Gaussian images as references. For the endoplasmic reticulum, the binary images were skeletonized. For mitochondria, the binary images were labeled by considering simply connected pixels as individuals. For the vacuole, the binary images were processed slice by slice over the z-stack. Each z plane was first skeletonized, then the background was flood filled, and the color of the slice was reversed so that the foreground represents the vacuole cross-section in the plane.

To get the statistics of organelles in different cells, we iterated over the cells in the segmented cell image for each field of view, then labeled and measured the area of each cell. Then we applied the cell mask to each organelle image across all z planes. For organelles other than the vacuole, the label and number of pixels of each label in the image are recorded. For the vacuole, we iterated over z planes, sought and recorded the disk with the largest cross-section area. The volumes of cells and vacuoles were estimated by treating the organism as cylinders with a height of their diameters. The non-organelle volumes were estimated by subtracting the six organelle volumes from the cellular volumes.

**5. Error Analysis of the Organelle Segmentation.** To analyze the organelle size error from the segmentation, we exported the "probability" provided by ilastik. For the foreground-background segmentation, pixels whose foreground probability exceeds 0.5 were identified as organelles. We listed all of the unique values in the probability tensor. The upper error pixels were those whose probability equaled the lowest value greater than 0.5, while the lower pixels were the highest value less than 0.5. For the peroxisome-vacuole images in the leucine experiment, the threshold was 0.33 for both channels. The ratios of upper and lower errors over the total volumes of different organelles are shown in Figure-S1-c.

**6. Calculation of Correlation Coefficient of Organelle Properties in the Glucose Experiment.** The organelle property data were grouped by the cells they belonged to. And the average volumes, total volumes, and counting numbers of the organelles inside each cell were calculated. These three statistics of the six organelles (except the counting number of the endoplasmic reticulum), along with the cellular statistics such as cross-section area, estimated volume, and characteristic length (estimated radii of the cross-section), were used to calculate the correlation coefficients between each pair of statistical variables.

**7. Log-log Regression Analysis of the Cell/Organelle Total Volume.** The organelle property data were aggregated by individual cells to get the total volume of each type of organelle. Then the cellular volumes, six organelle volumes, and non-organelle volumes were taken logarithm. For each experiment, the smallest 10% of the volumes were excluded to reduce the statistical fluctuation. We applied a linear regression analysis to each pair of logarithms and recorded the slope in an 8x8 matrix. We also plotted the logarithms in an 8x8 grid scatter plot, where different colors label the experiment conditions.

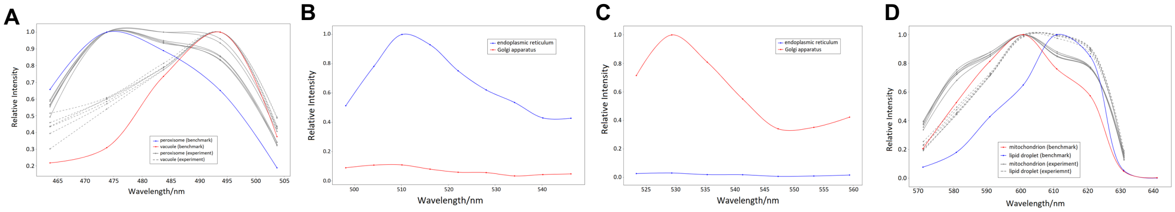


Figure S1. Experimentally measured fluorescence spectra used for spectral demixing. Spectra obtained from 445nm (A), 488nm (B), 514nm (C), and 561nm (D) excitation lasers. Colored lines indicate reference spectra obtained from the regions of interest from spectral images used to perform spectral demixing. Grey lines in (A,D) indicate spectra from pixels associated to each segmented organelle averaged over all pixels with individual grey lines corresponding to individual perturbations (i.e., glucose, leucine, $\beta$-estradiol, 1-nm-pp1, and rapamycin). Spectra in (A,D) are normalized with respect to their own maxima, while those in (B,C) are normalized by the maxima of the spectra of the target organelle, because the other organelles have near-zero signals. The avoided cross-talk in (B) and (C) support the use of the normal confocal rather than the hyperspectral imaging in the corresponding wavelength ranges.

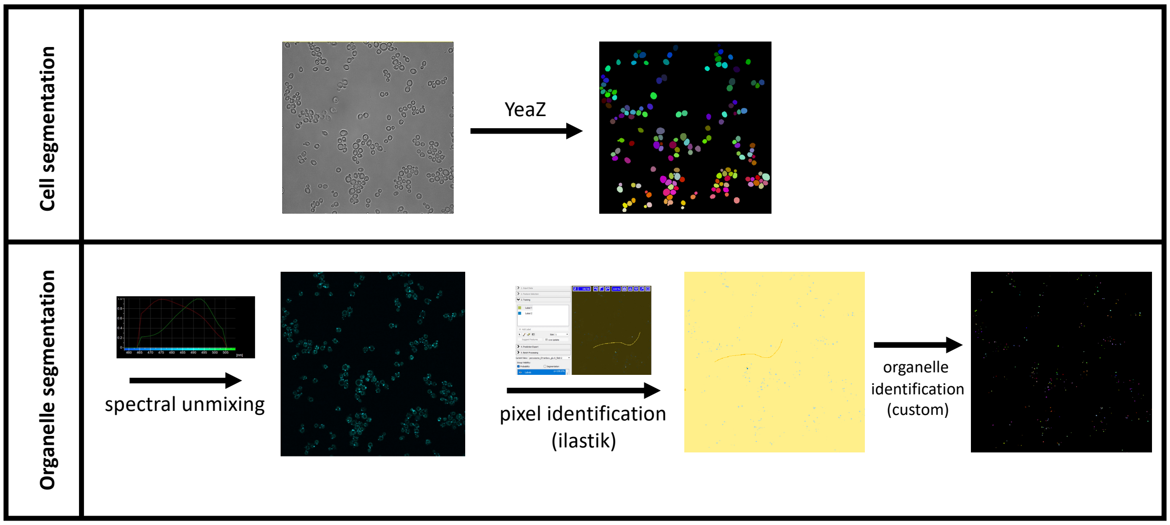


Figure S2. Schematic depicting the steps in image processing. Top row: brightfield cell image segmentation is performed with the YeaZ method (REF). Left panel: raw brightfield image. Right panel: Mask of segmented cells identified by YeaZ. Bottom row: fluorescent organelle image segmentation. The first step involves spectral demixing of hyperspectral image data into individual organelle channels (Fig. S1). The second step involves training the ilastik tool to distinguish between background and fluorescence-containing pixels for automated pixel classification. The third step involves grouping neighboring pixels into structures representing organelles.

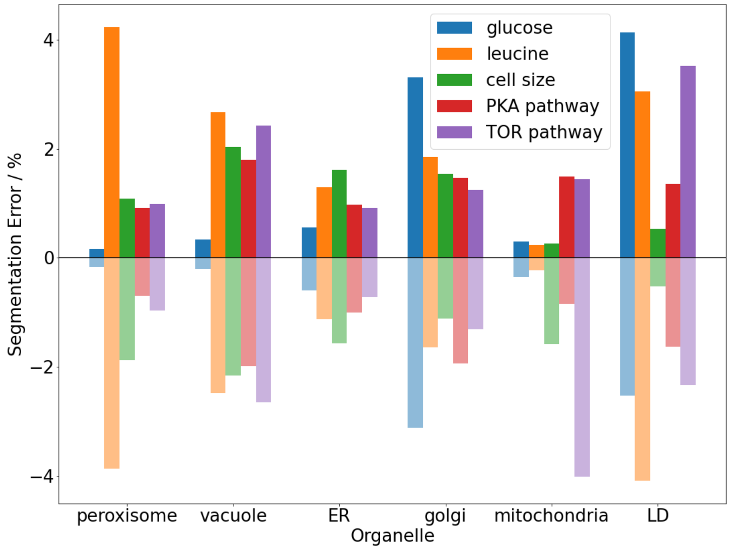


Figure S3. Result of error propagation from uncertainty in pixel classification to a given organelle to the error in the resulting volume fraction measurements.

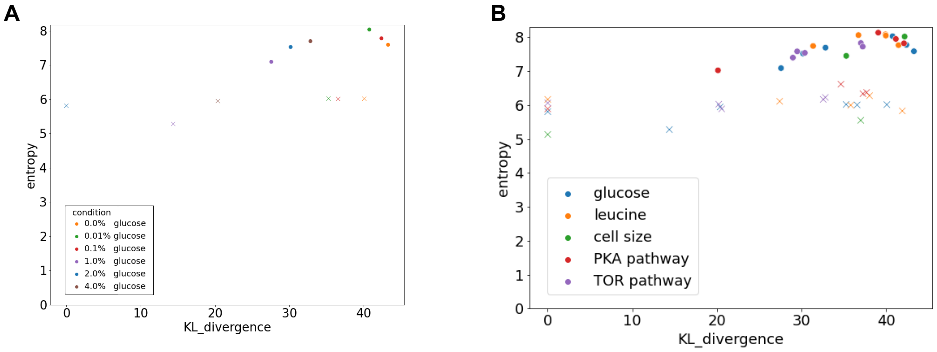


Figure S4. Mutual information between single cell organelle volume fraction joint probability distributions from indicated glucose concentration to the distribution from cells cultured in 2% glucose as a function of the Shannon entropy of the single cell organelle volume fraction joint probability distribution from indicated glucose concentration.

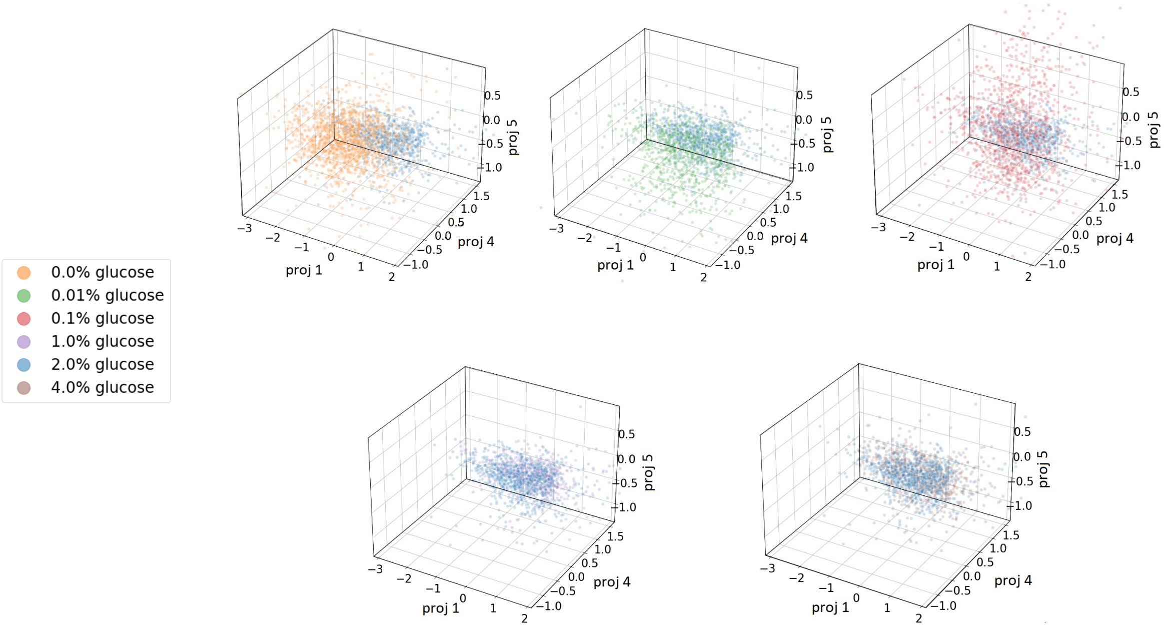


Figure S5. Single cell projections onto principal components obtained in cells grown in indicated glucose concentrations. The data from the cells here were used to compute the scalar products with the condition vector and the projection on the organelle volume fraction basis vectors depicted in Figure 2.

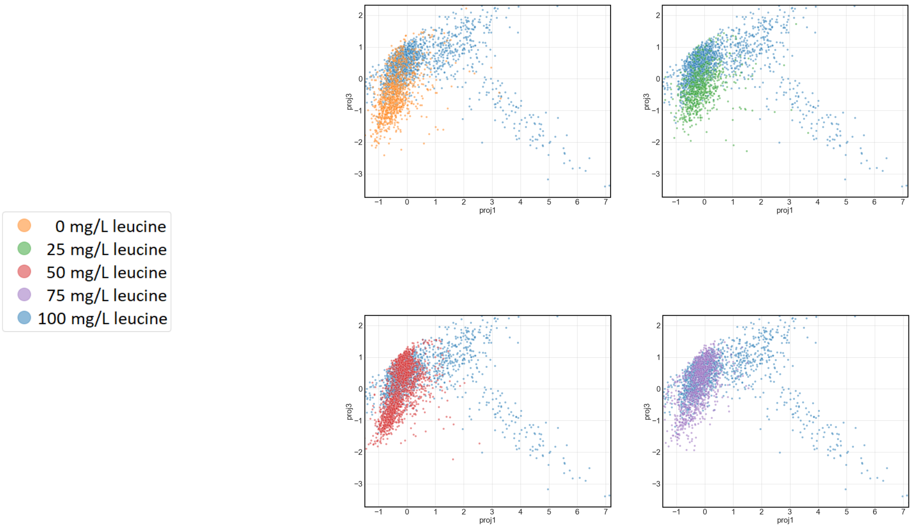


Figure S6. Single cell projections onto principal components obtained in cells grown in indicated leucine concentrations. The data from the cells here were used to compute the scalar products with the condition vector and the projection on the organelle volume fraction basis vectors depicted in Figure 3.

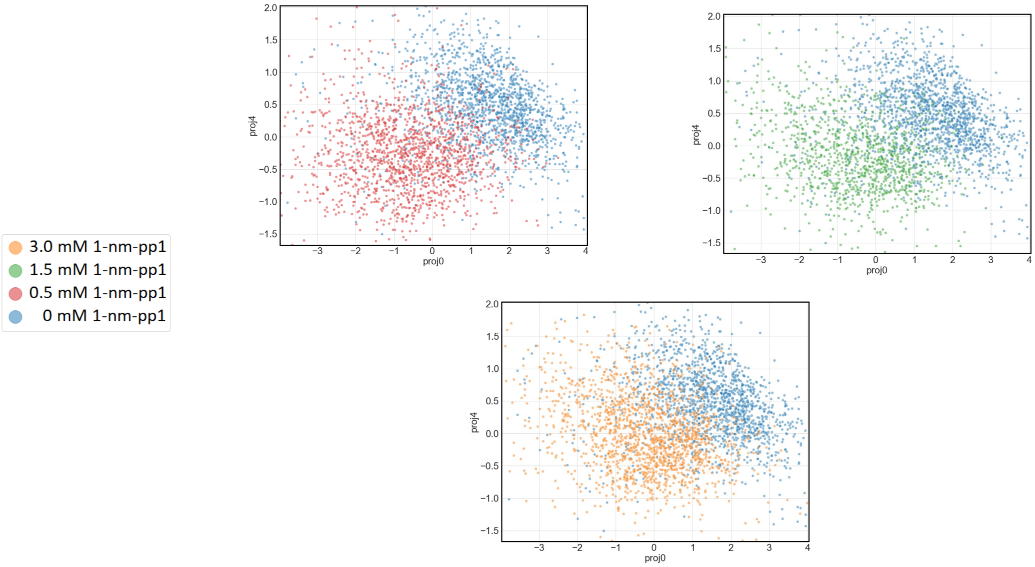


Figure S7. Single cell projections onto principal components obtained in cells grown in indicated 1-nm-pp1 concentrations to achieve a given PKA signaling activity level. The data from the cells here were used to compute the scalar products with the condition vector and the projection on the organelle volume fraction basis vectors depicted in Figure 3.

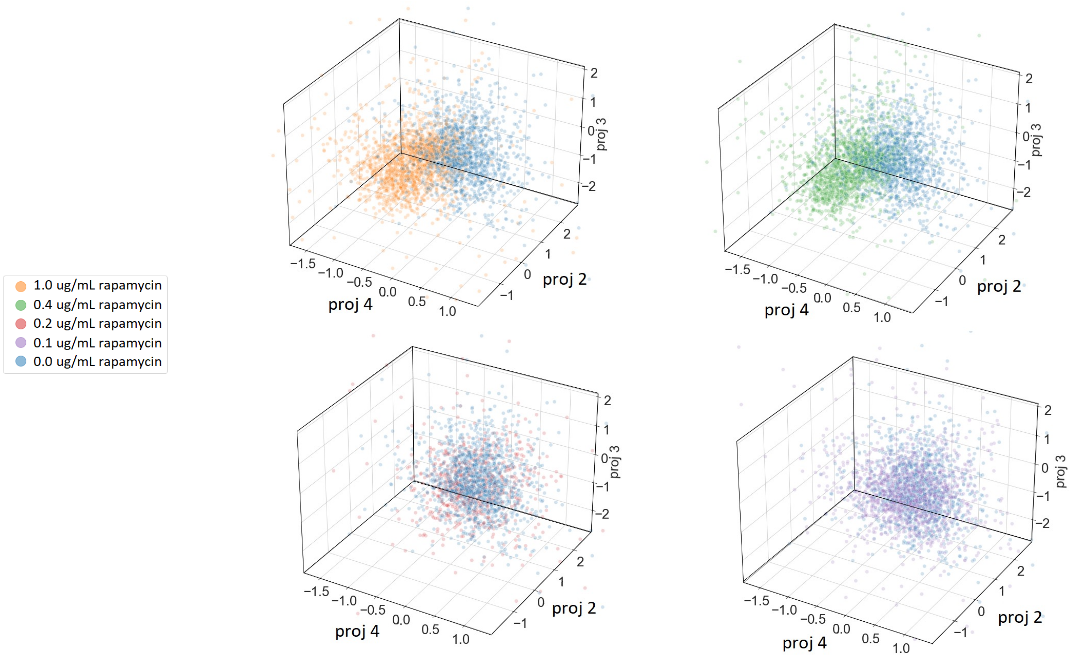


Figure S8. Single cell projections onto principal components obtained in cells grown in indicated rapamycin concentrations to achieve a given TOR signaling activity level. The data from the cells here were used to compute the scalar products with the condition vector and the projection on the organelle volume fraction basis vectors depicted in Figure 3.

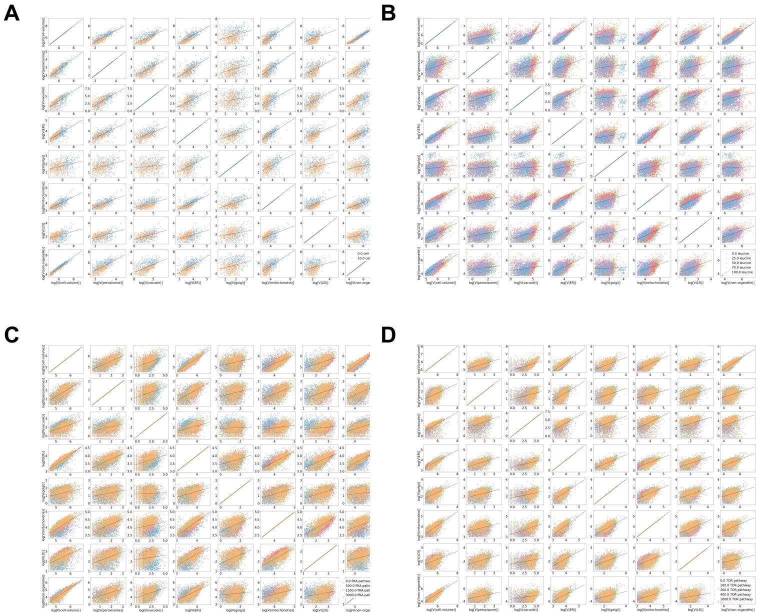


Figure S9. log-log volume scaling plots for cell size, leucine, PKA and TOR perturbations log-log organelle volume, cell volume, and nucleocytoplasmic volume plots shown from cells grown in indicated beta-estradiol, leucine, 1-nm-pp1, or rapamycin concentrations.

| | Cell Volume | Peroxisome | Vacuole | ER | Golgi | Mitochondrion | Lipid droplet | Nucleocytoplasm |
|---|---|---|---|---|---|---|---|---|
| Cell Volume | 1.000000 | 0.473681 | 0.240247 | 1.500602 | 0.596133 | 1.760848 | 0.513762 | 0.561543 |
| Peroxisome | 2.110921 | 1.000000 | 0.339950 | 2.908647 | 1.974009 | 3.459633 | 1.465634 | 0.806879 |
| Vacuole | 4.162221 | 2.941364 | 1.000000 | 6.851898 | 8.154160 | 6.910393 | 4.784052 | 4.007600 |
| ER | 0.666366 | 0.343787 | 0.145940 | 1.000000 | 0.469813 | 1.093088 | 0.446601 | 0.200564 |
| Golgi | 1.676972 | 0.506491 | 0.122620 | 2.128329 | 1.000000 | 4.359867 | 0.772015 | 0.352269 |
| Mitochondrion | 0.567846 | 0.289035 | 0.144710 | 0.914799 | 0.229307 | 1.000000 | 0.482187 | 0.100485 |
| Lipid droplet | 1.946039 | 0.682145 | 0.209004 | 2.239077 | 1.294744 | 2.073849 | 1.000000 | 0.090233 |
| Nucleocytoplasm | 1.780790 | 1.238844 | 0.249459 | 4.985924 | 2.837672 | 9.951608 | 11.07250 | 1.000000 |

Table S4. Volume scaling exponents derived from the slope of log-log volume plot in Fig. 1D

| | Cell Volume | Peroxisome | Vacuole | ER | Golgi | Mitochondrion | Lipid droplet | Nucleocytoplasm |
|---|---|---|---|---|---|---|---|---|
| Cell Volume | 1.000000 | 1.254028 | 0.338699 | 2.195181 | 1.489254 | 1.820387 | 1.894578 | 0.847974 |
| Peroxisome | 0.797405 | 1.000000 | 0.276740 | 1.734855 | 0.904057 | 1.433322 | 1.233213 | 0.611902 |
| Vacuole | 2.952522 | 3.613474 | 1.000000 | 6.809240 | 15.029861 | 6.733994 | 5.539209 | 2.916087 |
| ER | 0.455538 | 0.576402 | 0.146856 | 1.000000 | 0.480915 | 0.804762 | 0.547111 | 0.345070 |
| Golgi | 0.671278 | 1.105880 | 0.066528 | 2.079168 | 1.000000 | 1.835158 | 1.765996 | 0.403529 |
| Mitochondrion | 0.549328 | 0.697647 | 0.148500 | 1.242548 | 0.544868 | 1.000000 | 0.733693 | 0.407228 |
| Lipid droplet | 0.527753 | 0.810723 | 0.180528 | 1.827633 | 0.566061 | 1.362794 | 1.000000 | 0.280584 |
| Nucleocytoplasm | 1.179280 | 1.634193 | 0.342912 | 2.897880 | 2.477721 | 2.455561 | 3.563466 | 1.000000 |

Table S5. Volume scaling exponents derived from the slope of log-log volume plot in Fig. 9A

| | Cell Volume | Peroxisome | Vacuole | ER | Golgi | Mitochondrion | Lipid droplet | Nucleocytoplasm |
|---|---|---|---|---|---|---|---|---|
| Cell Volume | 1.000000 | 0.289917 | 0.219540 | 1.789634 | 0.120703 | 1.517627 | 1.347611 | 0.744620 |
| Peroxisome | 3.449038 | 1.000000 | 0.297917 | 6.770724 | 1.141342 | 5.842651 | 5.492304 | 3.292313 |
| Vacuole | 4.555053 | 3.356341 | 1.000000 | 8.261780 | 11.336064 | 6.375858 | 6.742737 | 9.013647 |
| ER | 0.558766 | 0.147692 | 0.121039 | 1.000000 | 0.096318 | 0.810337 | 0.674465 | 0.324782 |
| Golgi | 8.281675 | 0.869795 | 0.088194 | 10.382523 | 1.000000 | 4.935763 | 5.195691 | 10.764558 |
| Mitochondrion | 0.658898 | 0.171143 | 0.156839 | 1.234029 | 0.202595 | 1.000000 | 0.855540 | 0.348003 |
| Lipid droplet | 0.741999 | 0.182044 | 0.148308 | 1.482609 | 0.192450 | 1.168727 | 1.000000 | 0.423455 |
| Nucleocytoplasm | 1.342914 | 0.303626 | 0.110932 | 3.078943 | 0.092805 | 2.873191 | 2.361339 | 1.000000 |

Table S6. Volume scaling exponents derived from the slope of log-log volume plot in Fig. 9B

| | Cell Volume | Peroxisome | Vacuole | ER | Golgi | Mitochondrion | Lipid droplet | Nucleocytoplasm |
|---|---|---|---|---|---|---|---|---|
| Cell Volume | 1.000000 | 0.560906 | 0.214982 | 1.415210 | 0.328634 | 1.210427 | 0.556215 | 0.633907 |
| Peroxisome | 1.782744 | 1.000000 | 0.302969 | 2.653108 | 0.956058 | 2.364315 | 1.104060 | 0.967311 |
| Vacuole | 4.651420 | 3.300369 | 1.000000 | 8.208706 | 21.543395 | 11.561301 | 8.641595 | 2.905351 |
| ER | 0.706607 | 0.376912 | 0.121819 | 1.000000 | 0.243679 | 0.795849 | 0.408534 | 0.374285 |
| Golgi | 3.042174 | 1.045488 | 0.046402 | 4.103681 | 1.000000 | 2.661589 | 1.145614 | 0.937274 |
| Mitochondrion | 0.826118 | 0.422930 | 0.086492 | 1.256494 | 0.375698 | 1.000000 | 0.552329 | 0.312730 |
| Lipid droplet | 1.797674 | 0.905612 | 0.115702 | 2.447711 | 0.872763 | 1.810477 | 1.000000 | 0.727429 |
| Nucleocytoplasm | 1.577497 | 1.033624 | 0.344179 | 2.671667 | 1.062955 | 3.197467 | 1.374218 | 1.000000 |

Table S7. Volume scaling exponents derived from the slope of log-log volume plot in Fig. 9C

| | Cell Volume | Peroxisome | Vacuole | ER | Golgi | Mitochondrion | Lipid droplet | Nucleocytoplasm |
|---|---|---|---|---|---|---|---|---|
| Cell Volume | 1.000000 | 0.596982 | 0.213432 | 1.656827 | 0.977433 | 1.128271 | 0.548224 | 0.781345 |
| Peroxisome | 1.674921 | 1.000000 | 0.197777 | 2.843135 | 1.878723 | 2.456168 | 0.924864 | 1.135248 |
| Vacuole | 4.685340 | 5.055783 | 1.000000 | 8.320104 | 7.607112 | 5.905464 | 6.159622 | 7.727282 |
| ER | 0.603545 | 0.351704 | 0.120189 | 1.000000 | 0.568545 | 0.632428 | 0.338941 | 0.378466 |
| Golgi | 1.023014 | 0.532186 | 0.131446 | 1.758809 | 1.000000 | 1.131887 | 0.549609 | 0.583715 |
| Mitochondrion | 0.886252 | 0.407018 | 0.169338 | 1.581130 | 0.883383 | 1.000000 | 0.523870 | 0.545798 |
| Lipid droplet | 1.823746 | 1.080885 | 0.162332 | 2.950194 | 1.819334 | 1.908739 | 1.000000 | 1.321201 |
| Nucleocytoplasm | 1.279825 | 0.880587 | 0.129402 | 2.642094 | 1.712772 | 1.831944 | 0.756574 | 1.000000 |

Table S8. Volume scaling exponents derived from the slope of log-log volume plot in Fig. 9D
